# Analyzing the vast coronavirus literature with CoronaCentral

**DOI:** 10.1101/2020.12.21.423860

**Authors:** Jake Lever, Russ B. Altman

**Affiliations:** Department of Bioengineering, Stanford University, 443 Via Ortega, Stanford, CA 94305

## Abstract

The global SARS-CoV-2 pandemic has caused a surge in research exploring all aspects of the virus and its effects on human health. The overwhelming rate of publications means that human researchers are unable to keep abreast of the research.

To ameliorate this, we present the CoronaCentral resource which uses machine learning to process the research literature on SARS-CoV-2 along with articles on SARS-CoV and MERS-CoV. We break the literature down into useful categories and enable analysis of the contents, pace, and emphasis of research during the crisis. These categories cover therapeutics, forecasting as well as growing areas such as “Long Covid” and studies of inequality and misinformation. Using this data, we compare topics that appear in original research articles compared to commentaries and other article types. Finally, using Altmetric data, we identify the topics that have gained the most media attention.

This resource, available at https://coronacentral.ai, is updated multiple times per day and provides an easy-to-navigate system to find papers in different categories, focussing on different aspects of the virus along with currently trending articles.

## Background

The pandemic has led to the greatest surge in biomedical research on a single topic in documented history (Fig 1). This research is valuable both to current researchers working to understand the virus and also to future researchers as they examine the long term effects of the virus on different aspects of society. Unfortunately, the vast scale of the literature makes it challenging to evaluate. Machine learning systems should be employed to make it navigable by human researchers and to analyze patterns in it.

**Figure 1:**
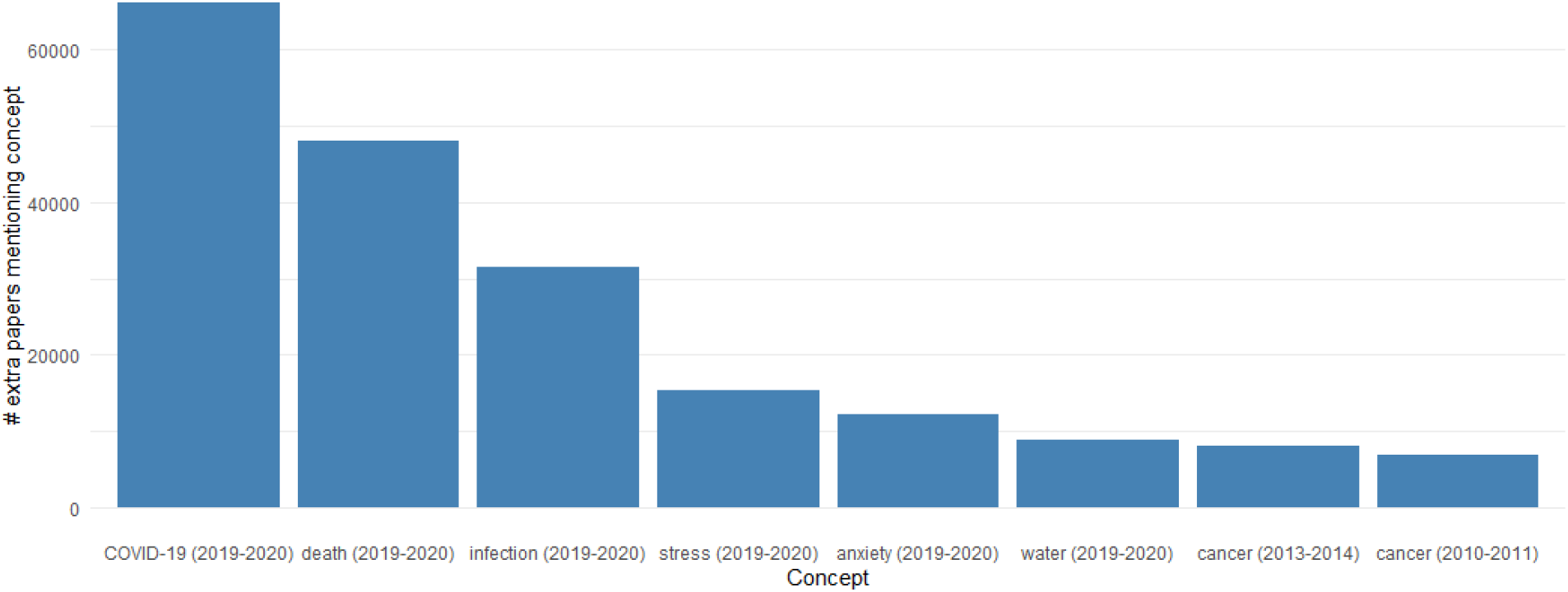
The change in focus of research on different biomedical concepts is measured using mentions of biomedical entities in PubTator. The greatest increase is seen by COVID research and unfortunately followed by death, infection, stress, and anxiety in the same time period.

Several methods have been built to make it easier to search and explore the coronavirus literature. LitCovid broadly categories the literature into 8 large categories, integrates with PubTator, and offers search functionality [1]. Collabovid uses the category data from LitCovid along with custom search functionality to provide another means of navigating the literature (accessible at https://www.collabovid.org). Other methods have developed different search interfaces to the literature such as Covidex [2]. Topic modeling approaches have also been employed to provide an unsupervised overview of major clusters of published articles [3] but are unable to provide the same quality as a supervised approach. COVID-SEE integrates several natural language processing analyses including search, unsupervised topic modeling, and word clouds [4]. The TREC-COVID shared task provided a number of information retrieval challenges on specific COVID-19 topics [5]. Apart from LitCovid’s limited set of categories, most approaches have avoided categorization and focussed on a search mechanism.

We present a detailed categorization system for coronavirus literature, integrated with search and esteem metrics to provide smooth navigation of the literature. We describe our efforts to maintain the CoronaCentral resource which currently categorizes 102,652 articles using machine learning systems based on manual curation of over 3000 articles and a custom category set. This work is designed to assist the research community in understanding the coronavirus literature and the continually-updated CoronaCentral dataset should help in analyzing a high-quality corpus of documents with cleaned metadata.

## Results

To provide more detailed and higher quality topics, we pursue a supervised learning approach and have annotated over 3,200 articles with categories from a set of 38 (Table 1). These categories cover the main topics of the papers (e.g. Therapeutics, Forecasting, etc) as well as specific article types (e.g. Review, Comment/Editorial, etc). Using a BERT-based document multi-label classification method, we achieved a micro-F1 score of 0.68 with micro-precision of 0.76 and micro-recall of 0.62. Table 2 provides a breakdown of the performance by category which shows varying quality of performance with some categories performing very well (e.g. contact tracing and forecasting) and others performing poorly (e.g. long haul) likely due to extremely low representation in the test set. Several other categories are identified using simple rule-based methods including the Book chapters, CDC Weekly Reports, Clinical Trials, and Retractions.

**Table 1:** Each article is labelled with multiple categories for their main topics and article type.

| Category | Description |
| --- | --- |
| Book Chapters | Book chapters that provide reviews or research on coronaviruses |
| CDC Weekly Reports | CDC Weekly Reports that provide a summary of the latest information about the pandemic |
| Clinical Reports | Clinical reports of patients discussing their symptoms or clinical course |
| Clinical Trials | Registered clinical trials for treating the virus |
| Comments & Editorials | Commentary, editorials and other short articles that are not original research |
| Communication | Research examining health messaging and communication with the general public and health professionals |
| Contact Tracing | Research studying the implementation and effect of contact tracing |
| Diagnostics | Research developing and testing methods to diagnose the virus |
| Drug Targets | Research discussing potential therapeutic coronavirus targets |
| Education | Research examining the effect of the coronavirus on the education system and how it has changed |
| Forecasting & Modelling | Research focussed on forecasting the spread or modelling the dynamics of the virus |
| Health Policy | Research discussing proposed and implemented health policy and its effect on the virus and society |
| Healthcare Workers | Research studying the effect of the virus on healthcare workers |
| Imaging | Research documenting CT and other scans of patients |
| Immunology | Research studying the interactions of the virus and the immune system |
| Inequality | Research studying the uneven health and economic outcomes for different groups |
| Infection Reports | Research documenting cases of virus spread through and between communities or countries |
| Long Haul | Research examining the long-term effects on coronavirus patients |
| Medical Devices | Research investigating medical devices used to treat the virus |
| Medical Specialties | Research examining the effect on and mitigations required by different medical specialties |
| Misinformation | Research studying the challenges of misinformation and deceptive communications involving the pandemic |
| Model Systems & Tools | Research proposing new model systems, tools and assays to study the virus |
| Molecular Biology | Research studying the molecular biology of the virus or host |
| News | News articles from academic journals and other sources indexed by PubMed or CORD-19 |
| Non-human | Research examining the effect of the virus in animals |
| Non-medical | Research examining the effect of the virus on other research areas (e.g. economics) |
| Pediatrics | Research studying the effect of the virus on children |
| Prevalence | Research studying the level of infections in different communities |
| Prevention | Research examining tools for slowing the spread of the virus (e.g. PPE) |
| Psychology | Research studying the effect on patients and healthcare workers mental health and understanding of risk |
| Recommendations | Research providing recommendations to medical professionals and others related to coronavirus |
| Retractions | Articles that have been withdrawn by the authors or the journal |
| Reviews | Review articles focussing on a variety of topics related to the virus |
| Risk Factors | Research examining risk factors for infection and severe symptoms |
| Surveillance | Research developing methods for surveillance of the disease spread |
| Therapeutics | Research studying the use of new and existing drugs for combating the virus |
| Transmission | Research studying the means of transmission of the virus |
| Vaccines | Research examining the development of vaccines |

**Table 2:** Machine learning results on different categories for held-out test set of 500 documents.

| Category | Precision | Recall | F1 |
| --- | --- | --- | --- |
| Clinical Reports | 0.84 | 0.63 | 0.72 |
| Comment/Editorial | 0.73 | 0.69 | 0.71 |
| Communication | 0.50 | 0.25 | 0.33 |
| Contact Tracing | 1.00 | 1.00 | 1.00 |
| Diagnostics | 0.93 | 0.83 | 0.88 |
| Drug Targets | 0.67 | 0.50 | 0.57 |
| Education | 0.63 | 0.71 | 0.67 |
| Effect on Medical Specialties | 0.79 | 0.49 | 0.61 |
| Forecasting & Modelling | 1.00 | 0.88 | 0.93 |
| Health Policy | 0.50 | 0.32 | 0.39 |
| Healthcare Workers | 0.80 | 0.89 | 0.84 |
| Imaging | 0.93 | 0.72 | 0.81 |
| Immunology | 0.44 | 0.44 | 0.44 |
| Inequality | 0.83 | 0.71 | 0.77 |
| Infection Reports | 0.75 | 0.30 | 0.43 |
| Long Haul | 0.00 | 0.00 | 0.00 |
| Medical Devices | 0.71 | 0.63 | 0.67 |
| Meta-analysis | 0.75 | 0.86 | 0.80 |
| Misinformation | 1.00 | 0.67 | 0.80 |
| Model Systems & Tools | 1.00 | 0.20 | 0.33 |
| Molecular Biology | 0.72 | 0.68 | 0.70 |
| News | 0.50 | 0.40 | 0.44 |
| Non-human | 0.83 | 0.50 | 0.63 |
| Non-medical | 1.00 | 0.43 | 0.61 |
| Pediatrics | 0.79 | 0.79 | 0.79 |
| Prevalence | 1.00 | 0.56 | 0.71 |
| Prevention | 0.72 | 0.56 | 0.63 |
| Psychology | 1.00 | 0.64 | 0.78 |
| Recommendations | 0.70 | 0.54 | 0.61 |
| Review | 0.57 | 0.75 | 0.65 |
| Risk Factors | 0.65 | 0.51 | 0.57 |
| Surveillance | 1.00 | 0.50 | 0.67 |
| Therapeutics | 0.70 | 0.68 | 0.69 |
| Transmission | 0.75 | 0.75 | 0.75 |
| Vaccines | 1.00 | 0.33 | 0.50 |
| MICRO | 0.76 | 0.62 | 0.68 |
| MACRO | 0.76 | 0.58 | 0.64 |

As of 21 December 2020, CoronaCentral covers 102,652 papers with Clinical Reports and the Effect on Medical Specialties being the most frequent categories (Fig 2). We made a specific effort to identify papers that discuss the effects on healthcare workers, the psychological aspects of the pandemic, the inequality that has been highlighted by the pandemic, and the long-term health effects of COVID. This final topic is covered by the Long Haul category which currently includes 239 papers. We find the first papers discussing the possible long-term consequences of COVID appeared in April 2020, for example, Kiekens et al [6]. Since then, there has been a slow steady increase in publications on the challenge of “Long COVID” with ∼20 papers per month recently. While all the annotated Long Haul documents used to train our system focus on SARS-CoV-2, our system finds 12 papers for the long-term consequences of SARS-CoV and one for MERS-CoV.

**Figure 2:**
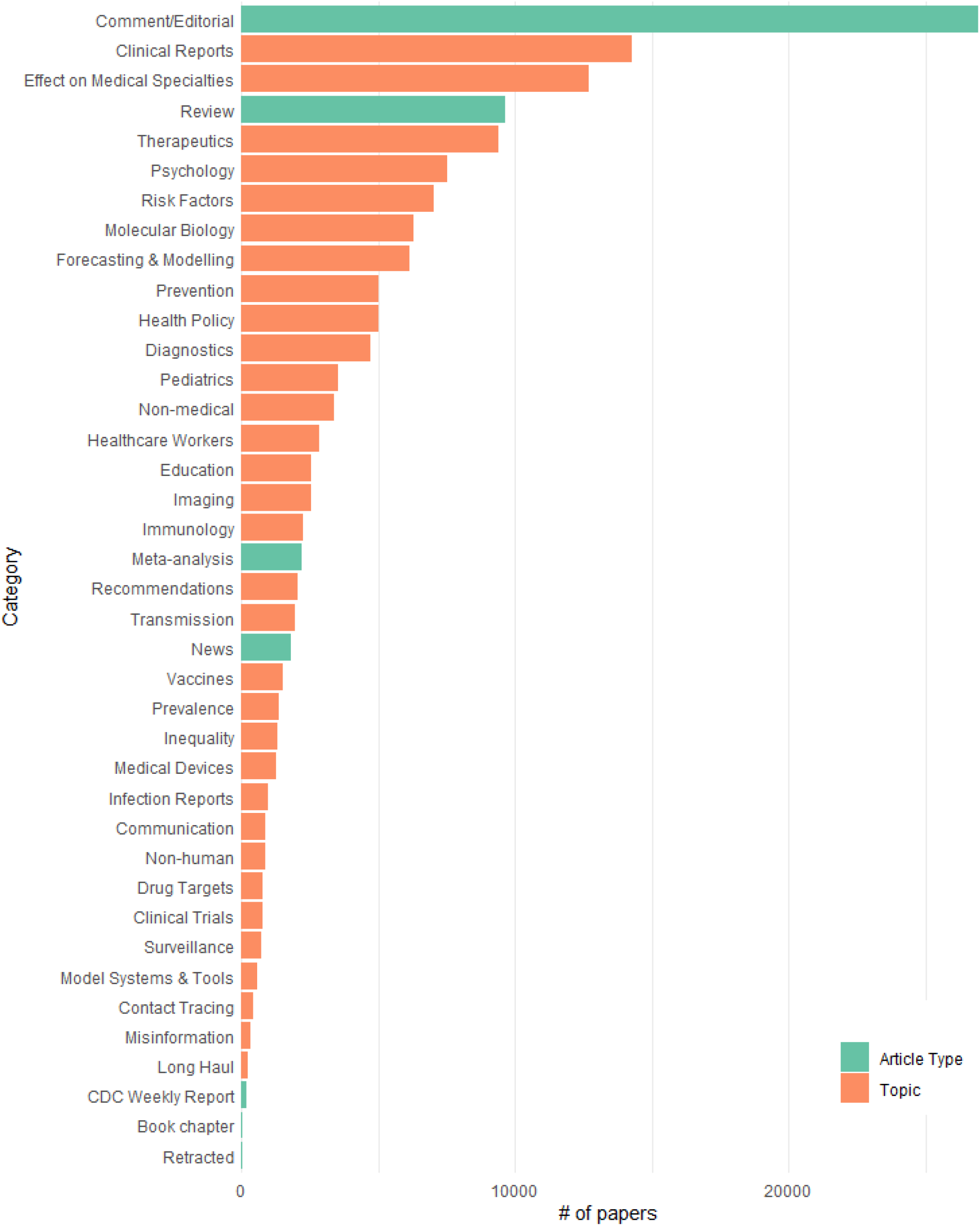
Frequency of each category across the entire coronavirus literature.

Identifying the type of publication is particularly important, given our estimate that 26.2% of coronavirus publications are comments or editorials and not original research. As well as the deep learning-based system for categorization, we use a web-crawling system to augment additional metadata including article type from PubMed and publishers’ websites. This automated categorization predicts papers as one of six types of article type, including Original Research, Meta-analysis, Review, Comment/Editorial, Book chapter, and News.

Figure 3 shows that different topics have drastically different distributions of article types. While almost all papers that look at forecasting or modeling the pandemic are original research, about half of the health policy articles are commentary or editorials. Notable topics with larger proportions of reviews are the more science-focused topics including Molecular Biology, Drug Targets, Therapeutics, and Vaccines. Clinical Trials and papers examining risk factors for coronavirus have a larger proportion of meta-analysis papers than other topics.

**Figure 3:**
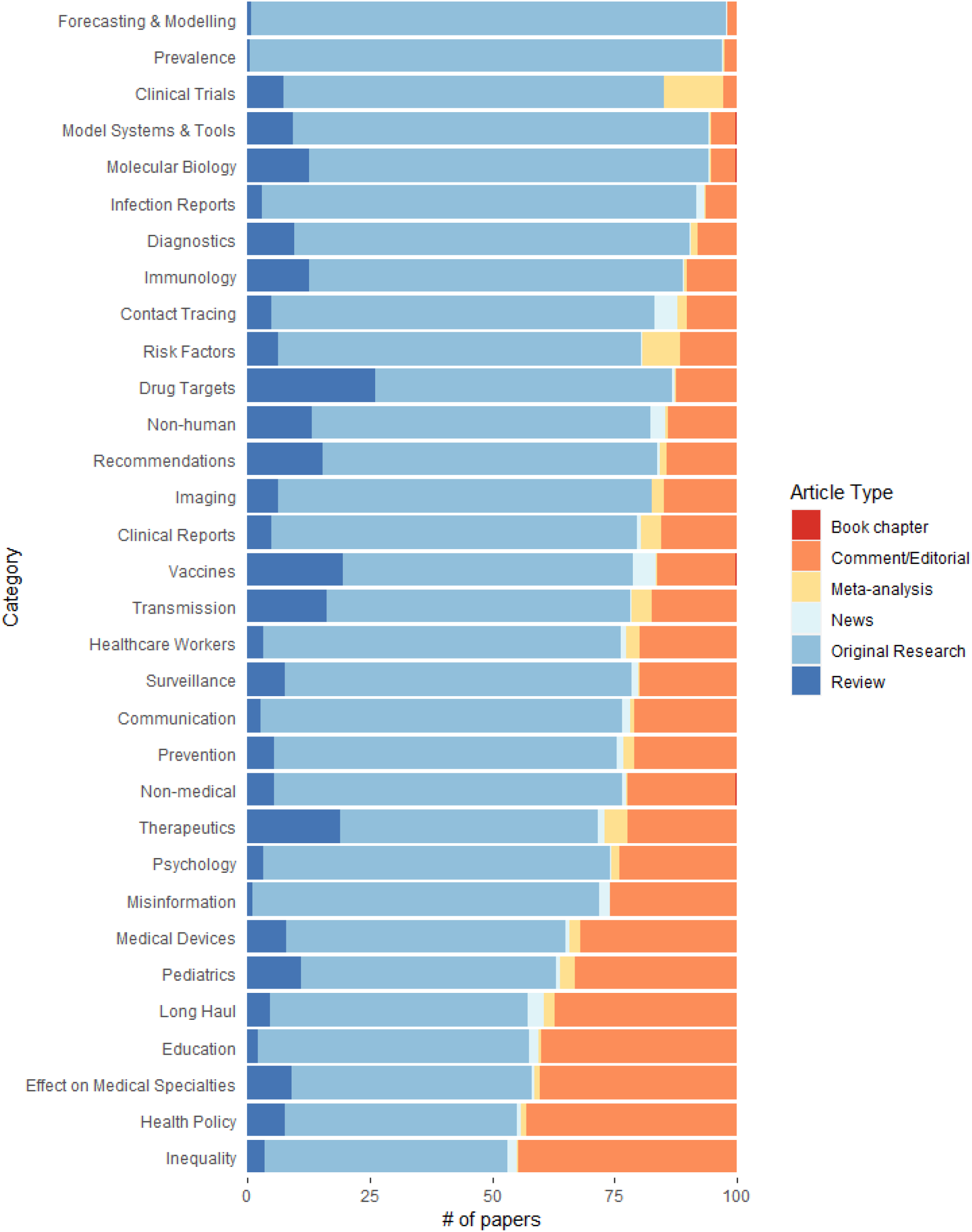
Different proportions of article types for each topic category

The predicted categories reveal the trend of publishing during the SARS-CoV-2 pandemic (Fig 4). Early original research focused on disease forecasting and modeling and has steadily decreased as a proportion compared to other areas of research, such as the risk factors of coronavirus, which have increased. Clinical reports that document patient symptoms have been steady, as a proportion, throughout the pandemic. In commentaries and editorials, the main topic has been the effect on different medical specialties (e.g. neonatology) and discussion on how the disciplines should adapt to the pandemic. Other common commentary topics include implementation of health policy and the psychological impact of the pandemic.

**Figure 4:**
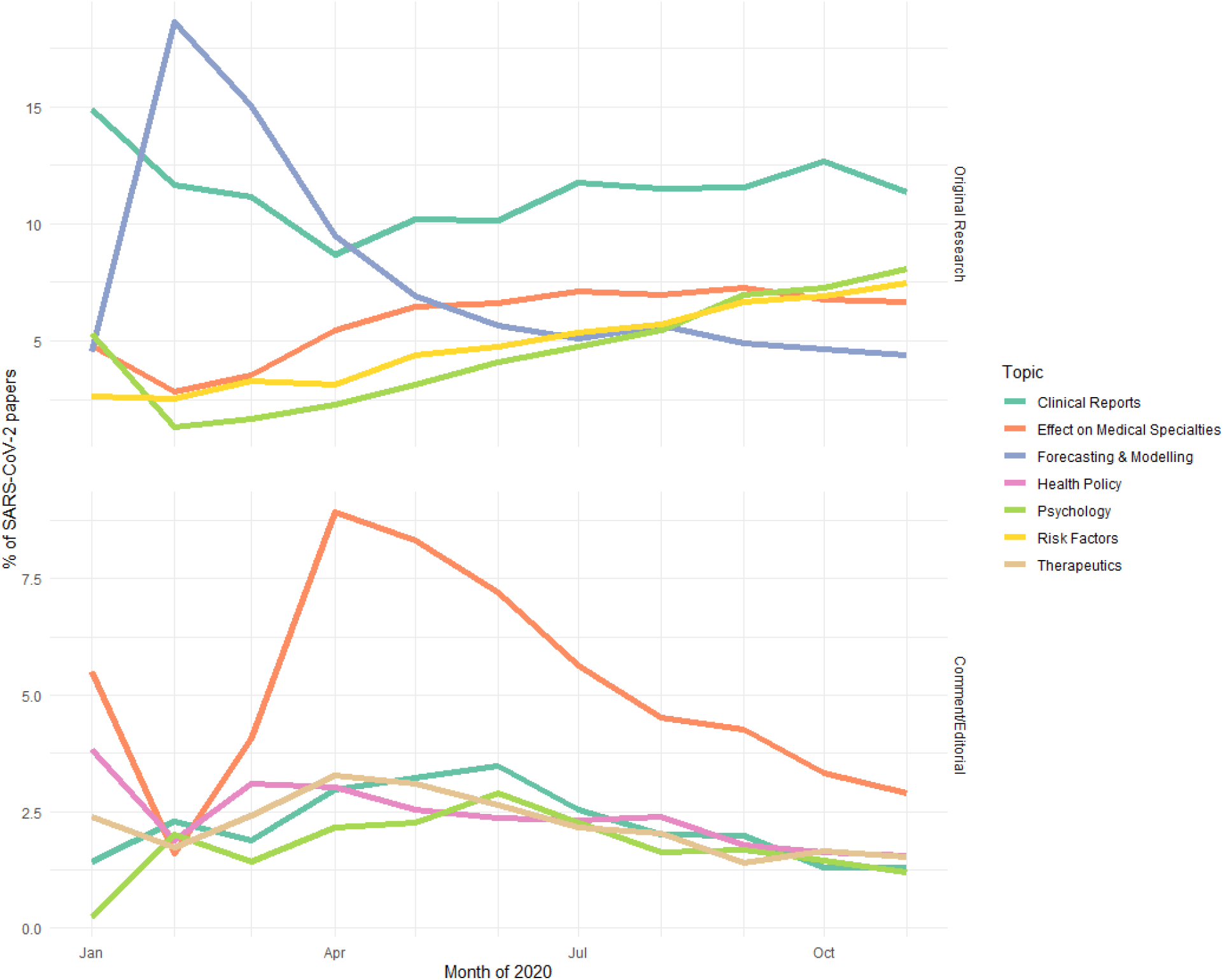
The trajectories of the top five topics for original research and comment/editorial articles for SARS-CoV-2.

Along with the article types and topics, we extract 13 relevant types of biomedical entities from the text to make the literature easier to navigate and identify important subtopics. Figure 4 provides a summary of the most common for each entity type broken down by the three coronaviruses. This includes geographic locations which enable quick identification of clinical reports in specific areas.

Preprint servers have proven incredibly important as Figure 6 shows with preprint servers leading the list of article sources. However, they only account for 5.7% of all articles. We find that the four indexed preprint servers have been used for dramatically different topics (Fig 7). As might be expected the more mathematically focussed papers, such as Forecasting/Modelling have been submitted to arXiv. Molecular biology research tends to go to bioRxiv and therapeutics research to ChemRxiv. MedRxiv has a more diverse clinical focused set of topics with the majority of the Risk Factors papers being sent there.

**Figure 5:**
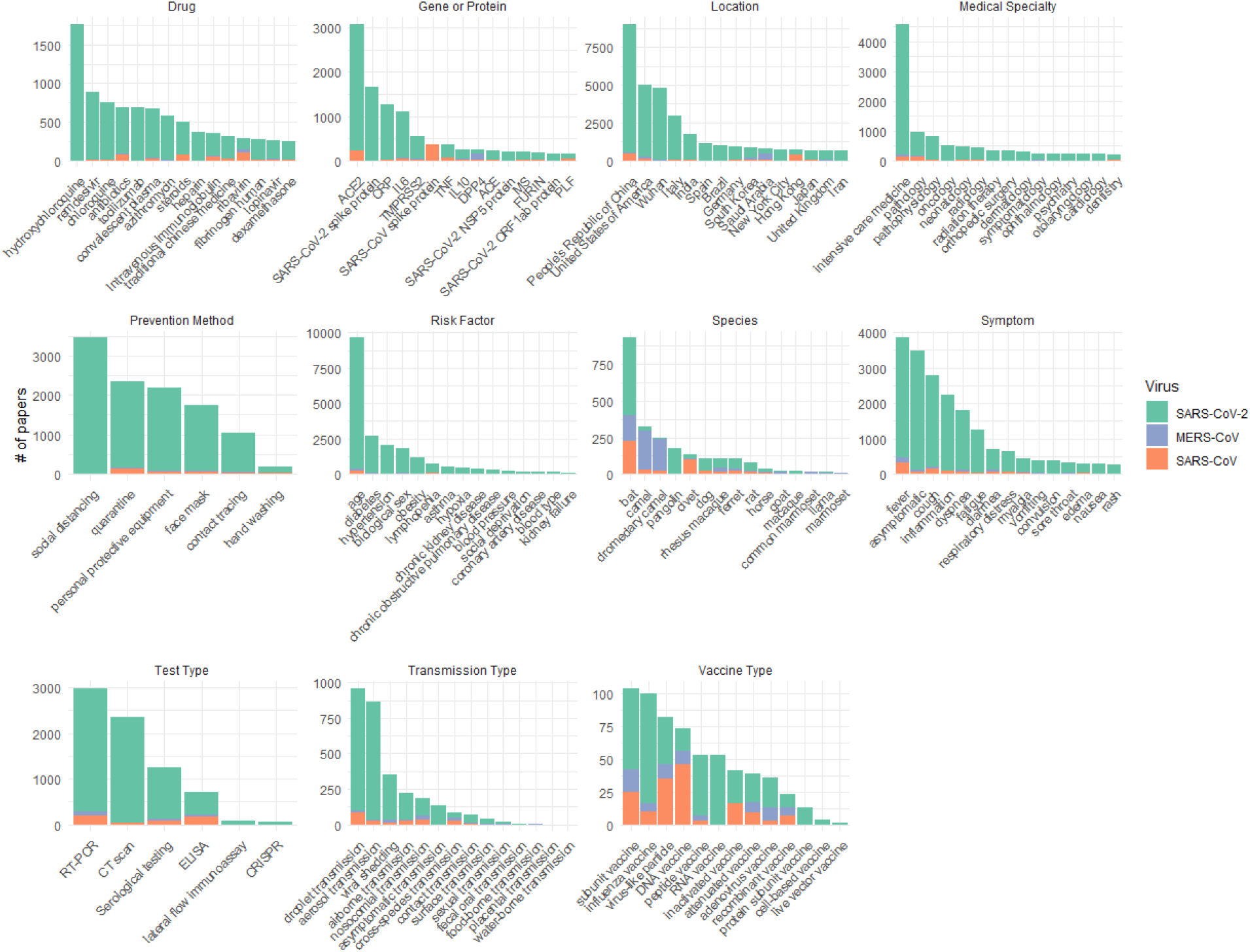
Top 15 entities for each entity type extracted from published literature for each virus.

**Figure 6:**
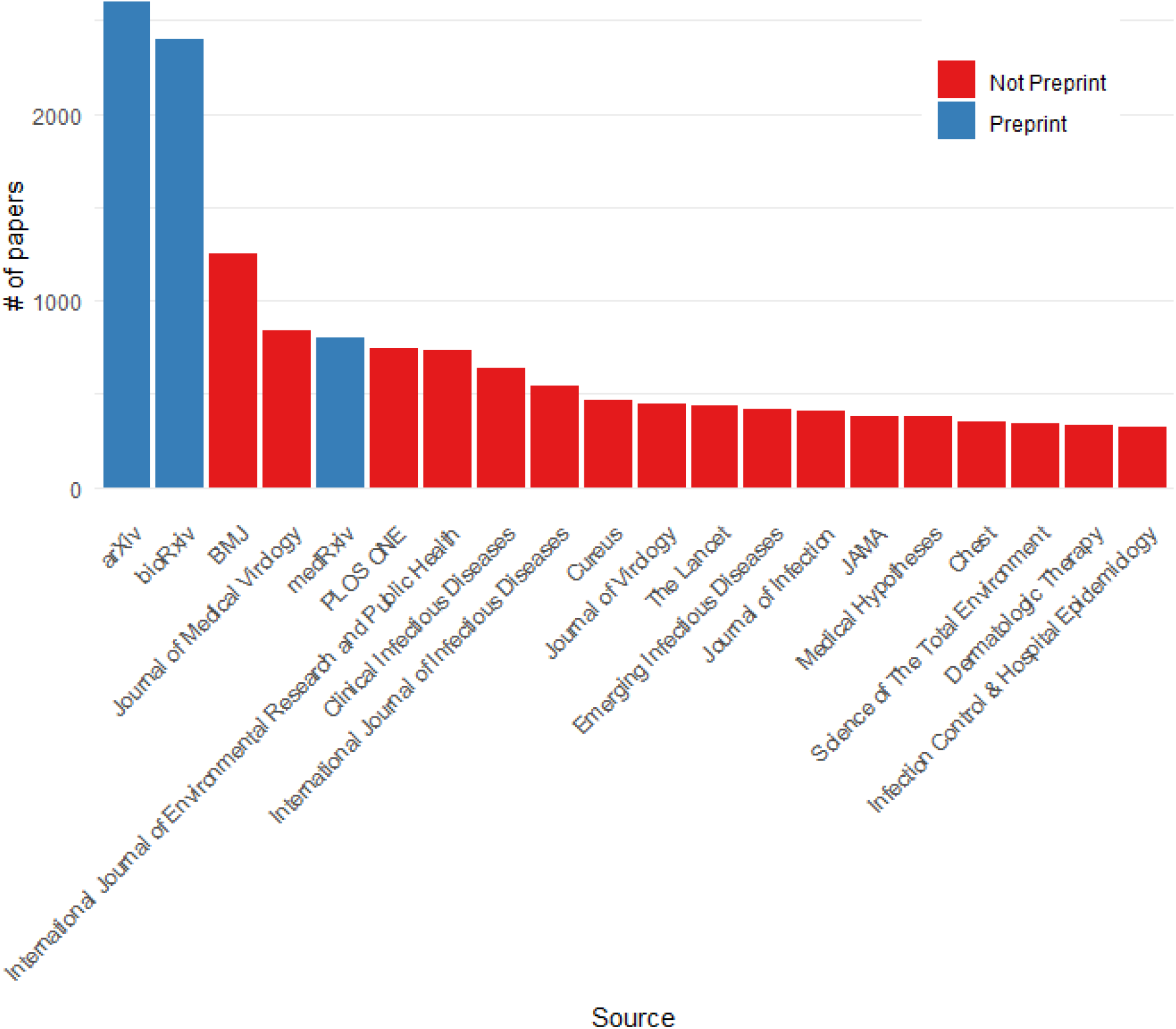
Top journals and preprint servers

**Figure 7:**
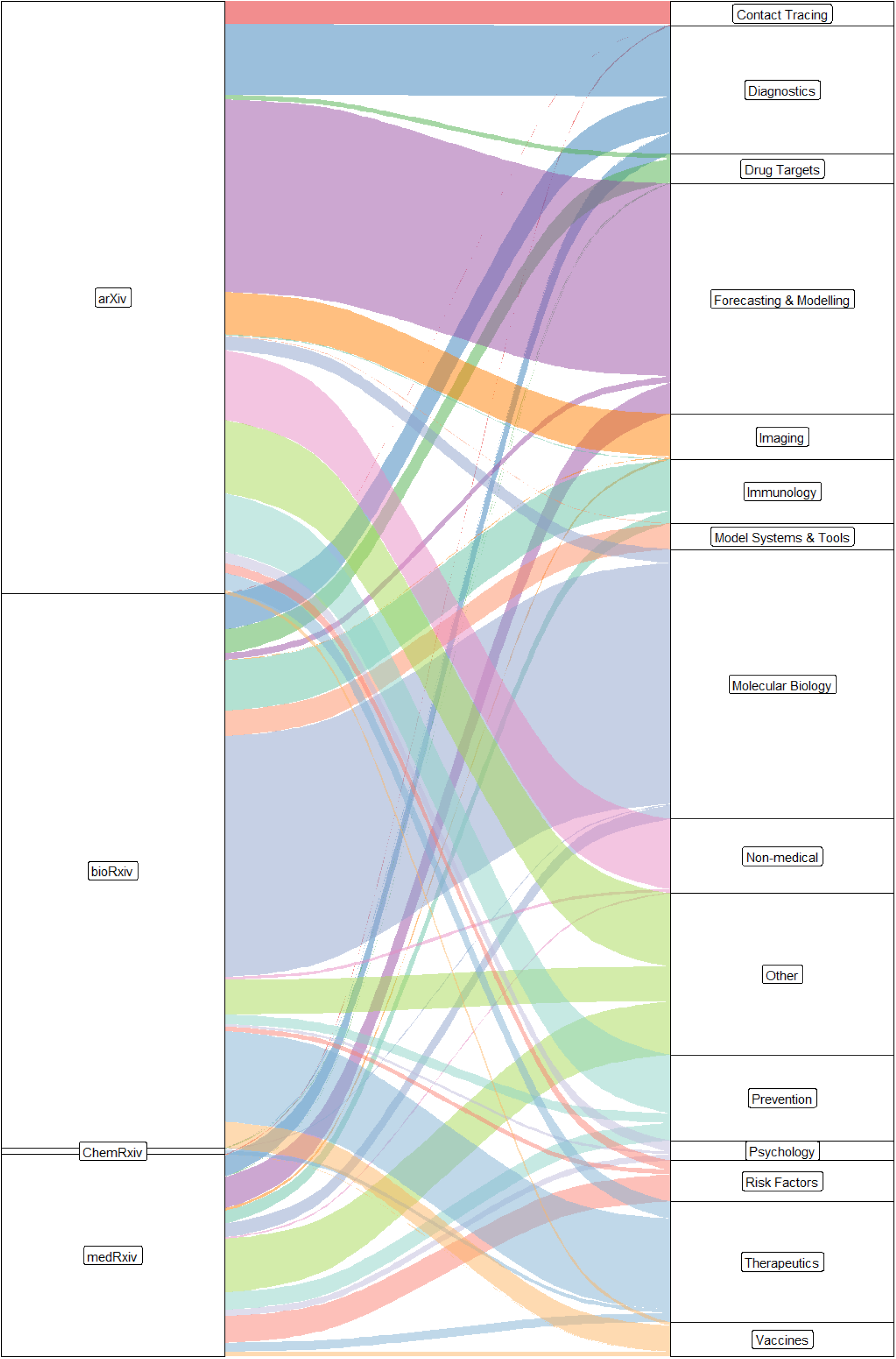
Topic breakdown for each preprint server and non-preprint peer-reviewed journals. Infrequent topics in preprints are grouped in Other.

The previous research on the SARS and MERS outbreaks are valuable sources of knowledge for viral biology, health policy implications, and many other aspects. We integrate research literature of these previous viruses along with SARS-CoV-2 and Figure 8 shows the different time ranges as well as the dramatic scale of the SARS-CoV-2 literature compared to the other two viruses. Notably, we are over the peak of SARS-CoV-2 literature, with 12,076 publications in May 2020. As an example of the strength of integrating previous coronavirus research, we identify drug candidates explored for SARS-CoV and MERS-CoV that have not yet appeared in SARS-CoV-2 publications (Table 3). Loperamide (Imodium) was found to inhibit MERS-CoV in low-micromolar concentrations in-vitro [7]. Two antibiotics (oritavancin and telavancin) were found to inhibit SARS and MERS viral entry and have not been further explored for SARS-CoV-2 [8].

**Figure 8:**
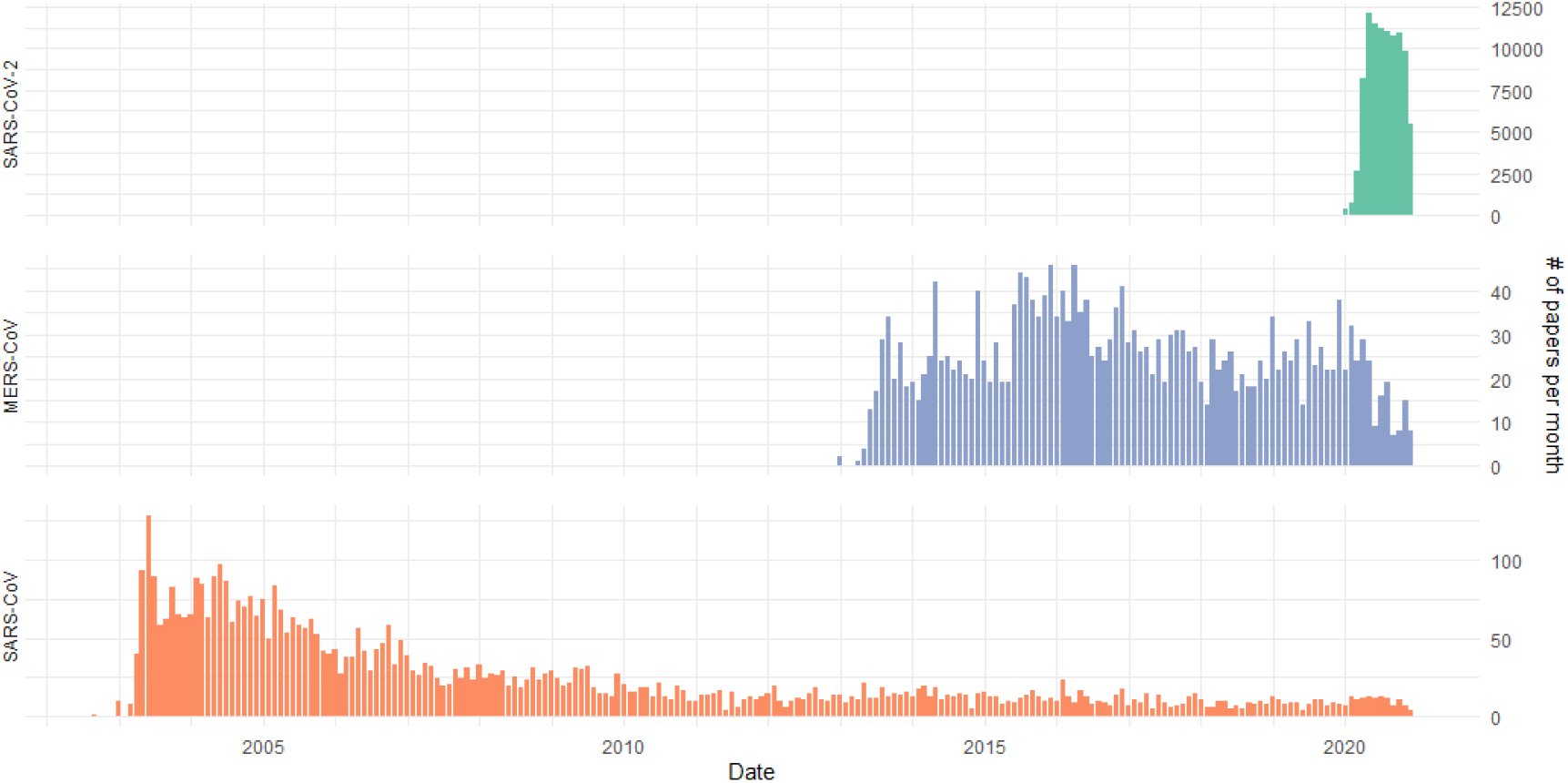
Publication rate for each virus

**Table 3:** Drugs discussed in SARS/MERS Therapeutics papers that have not appeared in SARS-CoV-2 papers

| Drug | # of SARS/MERS Papers | Effective in SARS/MERS screening | Description |
| --- | --- | --- | --- |
| enfuvirtide | 5 | No | HIV fusion inhibitor |
| Interferon alfacon-1 | 4 | Unclear | Interferons |
| hexachlorophene | 3 | Yes | Disinfectant |
| alendronic acid | 2 | Not relevant | Used for osteonecrosis |
| ethacrynic acid | 2 | Yes, for derivations | Loop diuretic |
| promazine | 2 | No | Antipsychotic |
| triclosan | 2 | No | Antibacterial/antifungal |
| Interferon Alfa-2a, Recombinant | 1 | Unclear, with ribavirin | Interferons |
| L-phenylalanine | 1 | Yes, for derivations | Isomer of amino acid |
| Nitroprusside | 1 | No | Blood pressure med |
| Palivizumab | 1 | Not relevant | Monoclonal antibody for RSV |
| adefovir dipivoxil | 1 | Not relevant | RTI for HepB |
| givosiran | 1 | Not relevant | Used for hepatic porphyria |
| loperamide | 1 | Yes | Diarrhea treatment |
| oritavancin | 1 | Yes | Antibiotic |
| phenazopyridine | 1 | Yes, for other coronaviruses | Analgesic |
| saracatinib | 1 | Yes | Kinase inhibitor |
| telavancin | 1 | Yes | Antibiotic |
| tobramycin | 1 | Not relevant | Antibiotic |

We integrate Altmetric data into CoronaCentral to identify papers that have received wide coverage in mass and social media. This enables users to quickly identify high-profile papers in each category as well as see currently trending articles. Figure 9 shows the breakdown of topics in the papers with the 100 papers with highest Altmetric scores. The distribution looks very different from the overall distribution of coronavirus literature with topics like Therapeutics, Transmission, and Prevention being more highly represented, reflecting the interest in understanding treatments and prevention methods.

**Figure 9:**
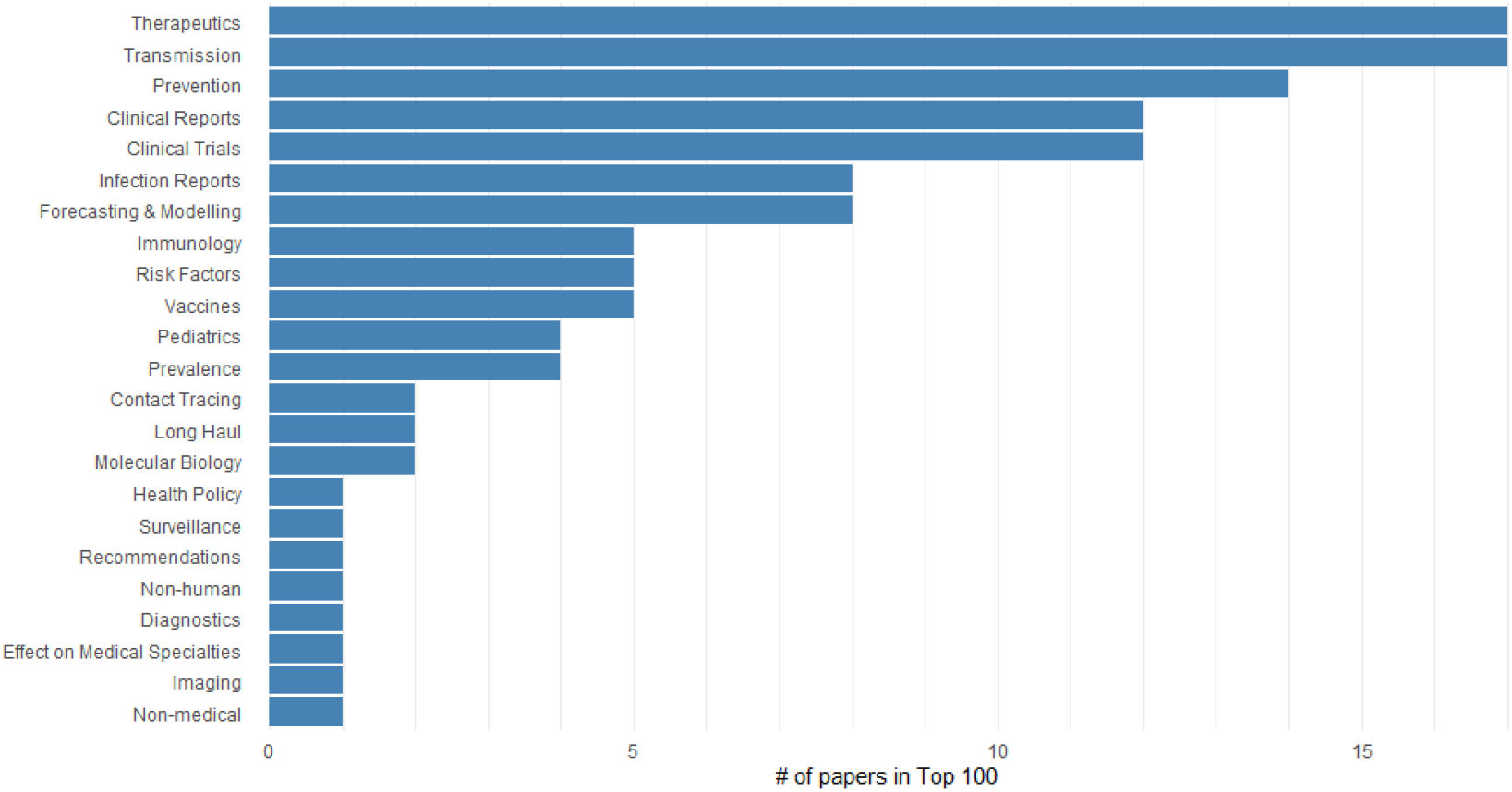
The number of papers categorized with each topic in the 100 papers with highest Altmetric scores

## Methods

### Data Collection

The CORD-19 dataset [9] and PubMed articles containing relevant coronavirus keywords are downloaded daily. Articles are cleaned to fix Unicode issues, remove erroneous text from abstracts, and identify publication dates. Non-English language articles are filtered out using a rule-based system based on sets of stopwords in multiple languages. To remove duplicates, documents were merged using identifiers, combinations of title and journal, and other metadata. Metadata from the publishers’ websites is also integrated which enables normalization of consistent journal names and further abstract text fixes. Additional manual fixes to title, abstracts, and metadata are applied to the corpus. Altmetric data is updated regularly and integrated with the data.

### Categories

Manual evaluation of an initial 1000 randomly selected articles was undertaken to produce a draft list of categories. These categories cover both the topic (e.g. Therapeutics) and the article type (e.g. Comment/Editorial). An iterative process was undertaken to adjust the category list to provide better coverage for the curated documents. A further 500 documents were sampled later in the pandemic and another iterative process was undertaken as new topics were appearing in larger quantities (e.g. contact tracing). Finally, several smaller topics that had not been captured by random sampling were identified and added to the category list (e.g. Long Haul). As the coronavirus literature grows, we may need to add new categories as new topics become more prominent.

### Category Annotation

Articles were manually annotated for categories using a custom web interface. The first 1500 randomly sampled articles were annotated during the iterative process that defined the set of categories. A further ∼1200 articles have been identified for annotation through manual identification, their high Altmetric scores or uncertainty in the machine learning system. Some of the articles were flagged using the CoronaCentral “Flag Mistake” system while others were identified through manual searching to improve representation of different topics. A final 500 articles were randomly selected and annotated for use as a held-out test set.

### Category Prediction

Cross-validation using a 75%/25% training/validation split was used to evaluate BERT-based document classifier as well as traditional methods as a baseline. Multi-label classifiers were implemented using ktrain [10] and HuggingFace models for BERT models and scikit-learn for others [11]. Hyperparameter optimization involved a grid search over parameters shown in Table 4 and selecting for the highest macro F1 score. The best model used the microsoft/BiomedNLP-PubMedBERT-base-uncased-abstract BERT model [12] with 32 epochs, a learning rate of 5e-05, and a batch size of 8. This model was then evaluated on the held-out test set for final performance and a full model was retrained using these parameters with all annotated documents and applied to the full coronavirus literature

**Table 4:** Parameter values searched for different classifier types

| Classifier | Parameter | Options |
| --- | --- | --- |
| BERT | Epochs | 4, 8, 12, 16, 24, 32, 48, 64, 80, 96 |
| BERT | Learning Rate | 1e-3, 5e-4, 1e-4, 5e-5, 1e-5, 5e-6 |
| BERT | Model | BlueBert, dmis-lab/biobert-v1.1, microsoft/BiomedNLP-PubMedBERT-base-uncased-abstract-fulltext, microsoft/BiomedNLP-PubMedBERT-base-uncased-abstract, allenai/scibert_scivocab_uncased, allenai/scibert_scivocab_cased' |
| Logistic Regression / Linear SVC / Random Forests | SVD Dimensionality Reduction | None, 8, 16, 32, 64, 128 |
| Logistic Regression / Linear SVC | C | 0.1, 1, 10, 20 |
| Random Forests | No of Estimators | 50, 100, 200 |

### Additional Category Identification

The BERT system predicts five article types but not Original Research. Documents are tagged as Original Research if not predicted as another article type. Clinical trials are identified through regular expression search for trial identifiers and book chapters for chapter headings in the title. The metadata provided by the publisher’s website is combined with PubMed metadata to identify some article types, e.g. documents tagged as Commentary or Viewpoints on publisher’s websites were categorized as Comment/Editorial. Retractions are identified through PubMed flags and titles beginning with “Retraction”, “Retracted” or “Withdrawn”.

### Entity Extraction

A set of entity types that would improve search across the literature was developed, e.g. drug names, locations, and more. This set was refined based on entities that would be particularly relevant for different categories (e.g. Drug for Therapeutics, Symptom for Clinical Reports, etc). A final 13 entity types were chosen and lists of entities were sourced from WikiData or built manually. Entities of types Drug, Location, Symptom, Medical Specialty, and Gene/Protein are gathered from Wikidata using a series of SPARQL queries. A custom list of Prevention Methods, Risk Factors, Test Types, Transmission Types, and Vaccine Types is also constructed based on Wikidata entities. Additional customizations are made to remove incorrect synonyms. A custom list of coronavirus proteins was added to the Gene/Protein list. Exact string matching is used to identify mentions of entities in text using the Wikidata set of synonyms and a custom set of stopwords. A simple disambiguation method was used to link virus proteins with the relevant virus based on mentions of the virus elsewhere in the document. This meant that a mention of a “Spike protein” in a MERS paper would correctly link it to the MERS-CoV spike protein and not to the SARS-CoV-2 spike protein. If multiple viruses were mentioned, no disambiguation was made.

### Interface

The data is presented through a website built using NextJS with a MySQL database backend. Visualizations are generated using ChartJS and mapping using Leaflet.

### PubTator Concept Analysis

To find the concepts that have had the largest difference in frequency, PubTator Central [13] was used as it covers a broad range of biomedical entity types such as disease, drug, and gene. It was aligned with PubMed and PubMed Central articles to link publication dates to entity annotations. This used the BioText project (https://github.com/jakelever/biotext). Concept counts were calculated per publication year and the differences between these ordered. Entity mentions of the type “Species” were removed due to lack of value as “human” dominated the data.

### Drug Analysis

To identifying drugs of interest from SARS and MERS research, SARS/MERS papers that were predicted to have the topic Therapeutics were filtered and drug mentions were extracted. These drug mentions were cross-referenced against all drug references in SARS-CoV-2 papers and those with a match were kept. The remaining drugs were manually reviewed using their source SARS/MERS papers to identify those that had shown efficacy in a SARS/MERS model.

### Other Analyses

All other analyses were implemented in Python and visualized using R and ggplot2.

### Code Availability

The code for the machine learning system and paper analysis are available at https://github.com/jakelever/corona-ml. The code for the web interface is available at https://github.com/jakelever/corona-web.

### Data Availability

The data is hosted on Zenodo and available at https://doi.org/10.5281/zenodo.4383289.

## Acknowledgements

This project has been supported by the Chan Zuckerberg Biohub and through the National Library of Medicine LM05652 grant (RBA).

## References

1. Chen Q, Allot A, Lu Z. Keep up with the latest coronavirus research. Nature. 2020;579: 193–193.

2. Zhang E, Gupta N, Tang R, Han X, Pradeep R, Lu K, et al. Covidex: Neural ranking models and keyword search infrastructure for the covid-19 open research dataset. arXiv preprint arXiv:200707846. 2020.

3. Doanvo A, Qian X, Ramjee D, Piontkivska H, Desai A, Majumder M. Machine learning maps research needs in covid-19 literature. Patterns. 2020; 100123. doi:https://doi.org/10.1016/j.patter.2020.100123

4. Verspoor K, Šuster S, Otmakhova Y, Mendis S, Zhai Z, Fang B, et al. COVID-see: Scientific evidence explorer for covid-19 related research. arXiv preprint arXiv:200807880. 2020.

5. Roberts K, Alam T, Bedrick S, Demner-Fushman D, Lo K, Soboroff I, et al. TREC-covid: Rationale and structure of an information retrieval shared task for covid-19. Journal of the American Medical Informatics Association. 2020.

6. Kiekens C, Boldrini P, Andreoli A, Avesani R, Gamna F, Grandi M, et al. Rehabilitation and respiratory management in the acute and early post-acute phase.“Instant paper from the field” on rehabilitation answers to the covid-19 emergency. Eur J Phys Rehabil Med. 2020; 06305–4.

7. De Wilde AH, Jochmans D, Posthuma CC, Zevenhoven-Dobbe JC, Van Nieuwkoop S, Bestebroer TM, et al. Screening of an fda-approved compound library identifies four small-molecule inhibitors of middle east respiratory syndrome coronavirus replication in cell culture. Antimicrobial agents and chemotherapy. 2014;58: 4875–4884.

8. Zhou N, Pan T, Zhang J, Li Q, Zhang X, Bai C, et al. Glycopeptide antibiotics potently inhibit cathepsin l in the late endosome/lysosome and block the entry of ebola virus, middle east respiratory syndrome coronavirus (mers-cov), and severe acute respiratory syndrome coronavirus (sars-cov). Journal of Biological Chemistry. 2016;291: 9218–9232.

9. Wang LL, Lo K, Chandrasekhar Y, Reas R, Yang J, Burdick D, et al. CORD-19: The COVID-19 open research dataset. Proceedings of the 1st workshop on NLP for COVID-19 at ACL 2020. Online: Association for Computational Linguistics; 2020. Available: https://www.aclweb.org/anthology/2020.nlpcovid19-acl.1

10. Maiya AS. Ktrain: A low-code library for augmented machine learning. arXiv. 2020;arXiv:2004.10703 [cs.LG].

11. Pedregosa F, Varoquaux G, Gramfort A, Michel V, Thirion B, Grisel O, et al. Scikit-learn: Machine learning in Python. Journal of Machine Learning Research. 2011;12: 2825–2830.

12. Gu Y, Tinn R, Cheng H, Lucas M, Usuyama N, Liu X, et al. Domain-specific language model pretraining for biomedical natural language processing. arXiv preprint arXiv:200715779. 2020.

13. Wei C-H, Allot A, Leaman R, Lu Z. PubTator central: Automated concept annotation for biomedical full text articles. Nucleic acids research. 2019;47: W587–W593.

